# Pre- and postsynaptically expressed spike-timing-dependent plasticity contribute differentially to neuronal learning

**DOI:** 10.1101/2021.09.01.458493

**Authors:** Beatriz E. P. Mizusaki, Sally S. Y. Li, Rui Ponte Costa, P. Jesper Sjöström

**Affiliations:** Centre for Research in Neuroscience, Brain Repair and Integrative Neuroscience Programme, Departments of Medicine, Neurology and Neurosurgery, The Research Institute of the McGill University Health Centre, Montreal General Hospital, Montreal, Quebec, Canada; Instituto de Física, Universidade Federal do Rio Grande do Sul, Porto Alegre, Rio Grande do Sul, Brazil; Computational Neuroscience Unit, Department of Computer Science, SCEEM, Faculty of Engineering, University of Bristol, Bristol, United Kingdom; Department of Physiology, University of Bern, Bern, Switzerland; Centre for Neural Circuits and Behaviour, Department of Physiology, Anatomy and Genetics, University of Oxford, Oxford, United Kingdom

## Abstract

A plethora of experimental studies have shown that long-term synaptic plasticity can be expressed pre- or postsynaptically depending on a range of factors such as developmental stage, synapse type, and activity patterns. The functional consequences of this diversity are not clear, although it is understood that whereas postsynaptic expression of plasticity predominantly affects synaptic response amplitude, presynaptic expression alters both synaptic response amplitude and short-term dynamics. In most models of neuronal learning, long-term synaptic plasticity is implemented as changes in connective weights. The consideration of long-term plasticity as a fixed change in amplitude corresponds more closely to post-than to presynaptic expression, which means theoretical outcomes based on this choice of implementation may have a postsynaptic bias. To explore the functional implications of the diversity of expression of long-term synaptic plasticity, we adapted a model of long-term plasticity, more specifically spike-timing-dependent plasticity (STDP), such that it was expressed either independently pre- or postsynaptically, or in a mixture of both ways. We compared pair-based standard STDP models and a biologically tuned triplet STDP model, and investigated the outcomes in a minimal setting, using two different learning schemes: in the first, inputs were triggered at different latencies, and in the second a subset of inputs were temporally correlated. We found that presynaptic changes adjusted the speed of learning, while postsynaptic expression was more efficient at regulating spike timing and frequency. When combining both expression loci, postsynaptic changes amplified the response range, while presynaptic plasticity allowed control over postsynaptic firing rates, potentially providing a form of activity homeostasis. Our findings highlight how the seemingly innocuous choice of implementing synaptic plasticity by single weight modification may unwittingly introduce a postsynaptic bias in modelling outcomes. We conclude that pre- and postsynaptically expressed plasticity are not interchangeable, but enable complimentary functions.

**Author summary:** Differences between functional properties of pre- or postsynaptically expressed long-term plasticity have not yet been explored in much detail. In this paper, we used minimalist models of STDP with different expression loci, in search of fundamental functional consequences. Biologically, presynaptic expression acts mostly on neurotransmitter release, thereby altering short-term synaptic dynamics, whereas postsynaptic expression affects mainly synaptic gain. We compared models where plasticity was expressed only presynaptically or postsynaptically, or in both ways. We found that postsynaptic plasticity had a bigger impact over response times, while both pre- and postsynaptic plasticity were similarly capable of detecting correlated inputs. A model with biologically tuned expression of plasticity also completed these tasks over a range of frequencies. Also, postsynaptic spiking frequency was not directly affected by presynaptic plasticity of short-term plasticity alone, however in combination with a postsynaptic component, it helped restrain positive feedback, contributing to activity homeostasis. In conclusion, expression locus may determine affinity for distinct coding schemes while also contributing to keep activity within bounds. Our findings highlight the importance of carefully implementing expression of plasticity in biological modelling, since the locus of expression may affect functional outcomes in simulations.

## Introduction

Learning and memory in the brain, as well as refinement of neuronal circuits and the development of receptive fields, are widely attributed to long-term synaptic plasticity [1]. While this notion is not yet formally experimentally proven [2], it has in recent years received strong experimental support in several brain regions, in particular the amygdala [3] and the cerebellum [4]. The notion that synaptic plasticity underlies memory is typically attributed to Hebb [5], but it is in actuality an idea that extends considerably farther back in time, e.g. to Ramon y Cajal and William James [6].

After the discovery by Bliss and Lømo [7] of the electrophysiological counterpart of Hebb’s postulate, now known as long-term potentiation (LTP), much effort has been focused on establishing the induction and expression mechanisms of long-term plasticity. In the 1990s, this led to a heated debate on the precise locus of expression of LTP, with some arguing for postsynaptic expression, whereas others were in favour of a presynaptic locus of LTP [8](Fig. 1A). Beginning in the early 2000’s, this controversy was gradually resolved by the realisation that plasticity depends critically on several factors, notably animal age, induction protocol, and precise brain region [9–11]. Indeed, this resolution has now been developed to the point that it is currently widely accepted that specific interneuron types have dramatically different forms of long-term plasticity [12, 13], meaning that long-term plasticity in fact depends on the particular synapse type [14]. In retrospect, it is probably not all that surprising that LTP in different circuits is expressed either pre- or postsynaptically, or both, given the diversity of computational functions of different synapses [15]. Nevertheless, the precise functional benefits of having LTP be expressed on one side of the synapse or the other have remained quite poorly explored, with only a handful of classical theoretical papers addressing this point [16–21].

**Fig 1.**
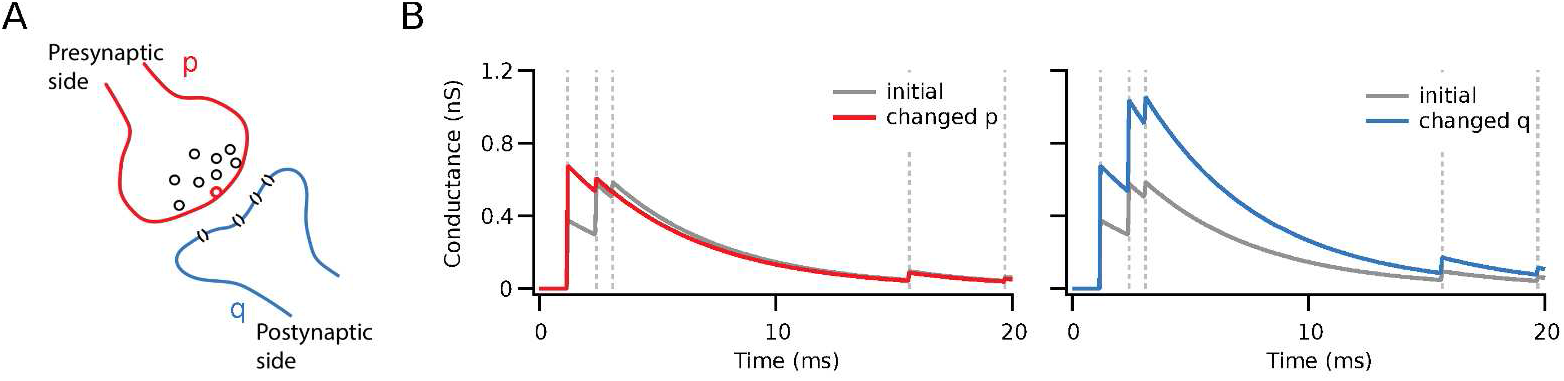
The postsynaptic response to the same stimulus after plasticity depends on the locus of expression. (**A**) Representation of pre- (red) and postsynaptic (blue) sides of a synapse, with probability of vesicle release *p*, and quantal amplitude *q*, i.e. the amplitude of postsynaptic response to a single vesicle. (**B**) Example of the difference between pre- and postsynaptic expression at inputs onto a cell. The identical initial response is illustrated in grey, while the potentiated responses are coloured red or blue. The amplitude of the first response after learning was set to be the same after pre- (red) and postsynaptic (blue) potentiation. With postsynaptic potentiation, the gain was increased by the same amount for all responses in the high-frequency burst. With presynaptic potentiation, however, the efficacy of the response train was redistributed toward its beginning, enhancing the first response but not the last.

Going back several decades, a multitude of highly influential computer models of neocortical learning and development have been proposed, some of them focusing on aspects such as the dependence of induction on firing rates [22–24], while others have emphasised the role of the relative millisecond timing of spikes in connected cells [25–27], and some yet have included both [28]. Irrespective of whether timing, rate, or other factors are used to determine the outcome of plasticity in theoretical models, it has virtually always been the case that – with a few notable exceptions [18, 19, 21] – the expression of plasticity itself has been regarded as a simple change in the magnitude of synaptic inputs between neurons of the network. As a minimal description this is of course a perfectly reasonable approach, as it is a parsimonious assumption that induction of long-term plasticity manifests itself in the alteration of connectivity weights.

However, the expression of plasticity is not always well modelled by this sole change of instantaneous magnitude. This is because presynaptically expressed plasticity leads to changes in synaptic dynamics, whereas postsynaptic expression does not (Fig. 1B). For instance, during high-frequency bursting, as the readily releasable pool of vesicles in a synaptic bouton runs out, leading to short-term depression of synaptic efficacy [29], while at other synapse types short-term facilitation dominates [30]. Such short-term plasticity is important from a functional point of view because it acts as a filter of the information that is transmitted by a synapse [31–33]. Short-term depressing connections are more likely to elicit postsynaptic spikes due to brief non-sustained epochs of activity, whereas facilitating synapses require that presynaptic activity be sustained for some period of time to elicit postsynaptic spikes. In other words, short-term facilitating connections act as high-pass filtering burst detectors [34, 35], while short-term depression provides low-pass filtering inputs more suitable for correlation detection and automatic gain-control [36–38]. As a corollary, it follows that presynaptic expression of plasticity may change the computational properties of a given synaptic connection. In this case, increasing the probability of release by the induction of LTP led to more prominent short-term depression due to depletion of the readily-releasable pool depletion, and as a consequence to a bias towards correlation detection at the expense of burst detection [39, 40].

Experimentally, it is long known that the induction of neocortical long-term plasticity may alter short-term depression [16, 41]. Although the functional consequences of short-term plasticity itself are as outlined above quite well described [39, 42], the theoretical implications of *changes* in short-term plasticity due to the induction of long-term plasticity are not well described. Yet, a majority of theoretical studies of long-term plasticity assumes that synaptic amplitude but not synaptic dynamics are altered by synaptic learning rules. One of the motivations of our present study is the observation that this seemingly innocuous assumption may not be neutral, but may in effect introduce a bias, because changing synaptic weight in theoretical models of long-term plasticity is equivalent to assuming that synaptic plasticity is solely postsynaptically expressed. This begs the question: What are the functional implications of pre- versus postsynaptically expressed long-term plasticity? Providing answers to this central issue is important for understanding brain functioning, as well as for knowing when weight-only changes in computer modelling is warranted, and when it is not.

Here, we use computational modelling to explore and describe the consequences of expressing plasticity pre- or postsynaptically in a single neuron under two simple paradigms (Fig. 2). One paradigm explores the postsynaptic response in relation to a repeated time-locked stimulus [26, 43, 44], while the other investigates the neuron’s ability to detect a correlated stimulus [45–47]. Initially, we compare and contrast relatively artificial scenarios, for which the locus of expression is either solely presynaptic, solely postsynaptic, or equally divided between both sides. We then move on to investigating the functional impact in a biologically realistic model with separate pre- and postsynaptic components that were tuned to experimental data from connections between neocortical layer-5 pyramidal cells [19]. We report that presynaptically expressed plasticity adjust the speed of learning, while postsynaptic expression is more efficient at regulating spike timing and frequency. We conclude that pre- and postsynaptically expressed plasticity enable different complimentary functions and are not equivalent.

**Fig 2.**
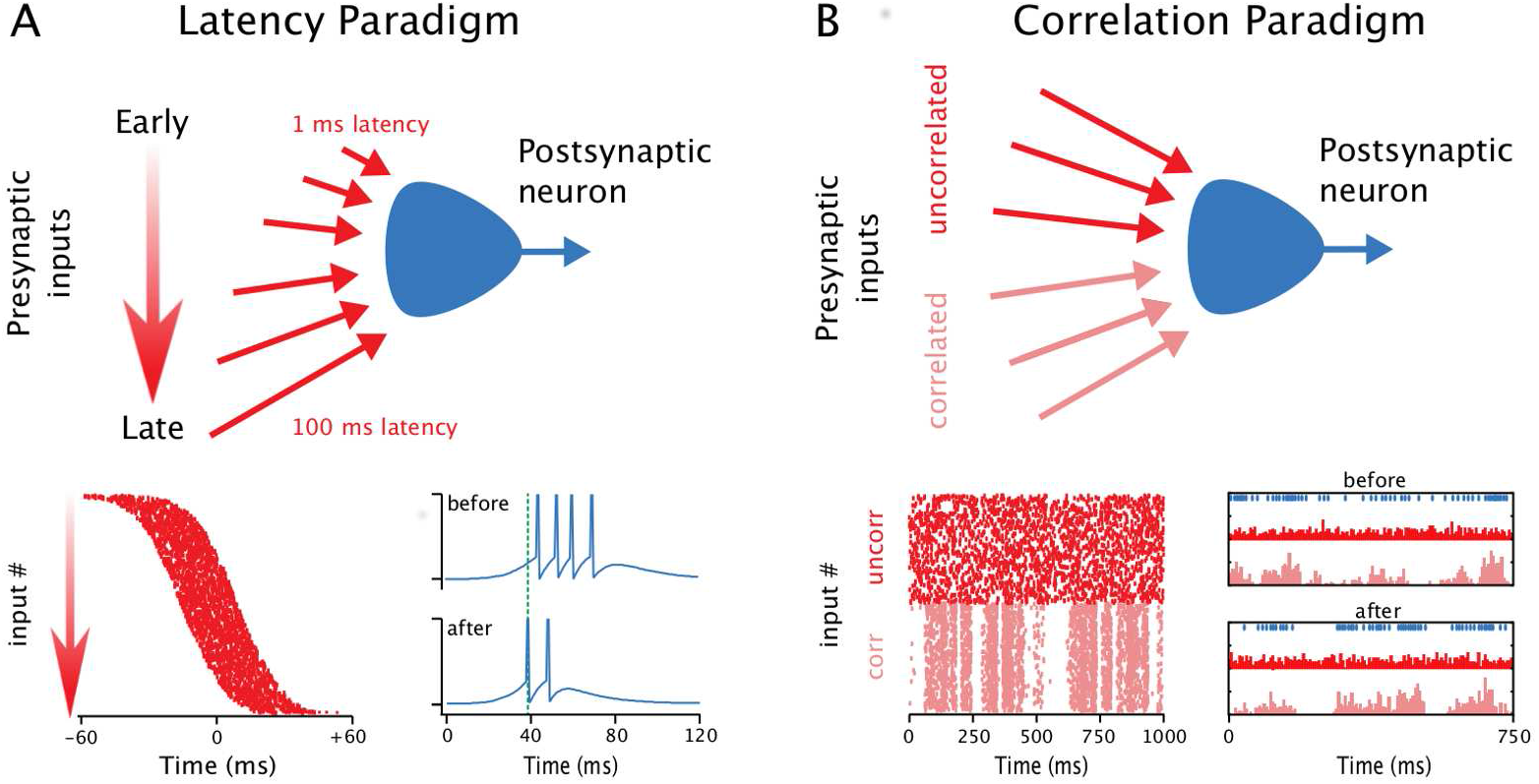
Two different STDP learning paradigms were explored. (**A**) Inputs arriving with a gradient of early to late timings resulted in reduced latency of the postsynaptic spiking response after STDP, as previously described [26]. In each trial, the postsynaptic neuron repeatedly received a brief volley of stimuli, between which short-term plasticity variables were allowed to return to their initial resting values. Bottom, left: Each presynaptic spike (raster dots) arrived with a different delay in the volley. Bottom, right: After a period of learning, the postsynaptic spiking response (blue) was shortened and started earlier. The learning task is thus to reduce the latency and to shorten the duration of the postsynaptic spiking response [26]. (**B**) Correlated inputs were selectively potentiated by STDP, as previously described [45]: The postsynaptic neuron received persistent stimulation, with half of the inputs having correlated activity, while the rest were uncorrelated. Bottom, left: Raster plot illustrating the correlated (corr) and uncorrelated (uncorr) input spiking. Bottom, right: After learning, the postsynaptic spiking (blue raster at top) was more correlated with the correlated inputs (pink histograms) than with the uncorrelated inputs (red histograms), indicating that the former drove postsynaptic activity. The learning task is thus to select for inputs that are correlated at the expense of those that are not [45]. In both paradigms, STDP is modelled with the same parameters, to permit comparison (e.g. timing window *τ* = 20 ms, see Methods).

## Results

Postsynaptically expressed plasticity is readily implemented as a simple change in synaptic gain, by adjusting the quantal amplitude, *q*. The impact of postsynaptic expression is therefore relatively unambiguous, since it scales all postsynaptic responses the same way. For example, in the case of repeated measures of presynaptic stimulation, the standard deviation and the mean of synaptic responses scale the same, so the coefficient of variation remains the same [48], which means synaptic noise levels remain the same after postsynaptically expressed plasticity.

Presynaptic plasticity, however, has at least two different distinct types of impact on a synapse. First, in the reliability of transmission due to stochastic vesicle release. Assuming release is binomially distributed, increasing the probability of release, *p*, increases the mean of synaptic responses while keeping the standard deviation roughly the same, which means the coefficient of variation is decreased after presynaptic LTP [48]. Second, increasing the probability of release depletes the pool o readily releasable pool of vesicles more rapidly. Therefore, synaptic short-term dynamics are necessarily changed by presynaptically expressed long-term plasticity, resulting in functional differences.

To limit the scope of the study, we focus on early forms of plasticity for which we have detailed experimental data [19]. We thus do not consider the possibility that the number of release sites, *n*, may change, as it does in late, protein-synthesis dependent forms of plasticity [49].

To distinguish between the two distinct types of impact of presynaptic plasticity, we model them separately. We start with presynaptic expression modelled as direct changes in the probability of vesicle release and compare that to postsynaptic expression. Subsequent to that, we model presynaptic expression as changes in short-term plasticity and compare that postsynaptic expression. This way, we aim to systematically tease apart any different contributions of the two distinct impacts of presynaptically expressed plasticity.

### Presynaptic expression modelled as changes in stochastic release

For the first set of simulations, we considered the probability of release (*p*_*j*_) and the quantal amplitude (*q*_*j*_), i.e. pre- and postsynaptic quantities, respectively. In each simulation, plasticity was expressed exclusively presynaptically, exclusively postsynaptically, or equally divided between both sides at connections onto a single-compartment point neuron (see Methods). Here, changes in *p*_*j*_ were explored in terms of their impact on stochastic release.

In the latency paradigm, in which a volley of stimuli arrives at the postsynaptic neuron with varying delays (Fig. 2A), plasticity resulted in the shortening of the time to respond — the latency — of the postsynaptic neuron, as well as a temporal sharpening of the response, with fewer spikes and shorter inter-spike intervals [26]. The average latency reduction (Fig. 3A, B), as well as the overall distribution of synaptic weights, decrease of postsynaptic activity duration and increase of postsynaptic firing frequency (Figs. 3C, D, E) did not differ appreciably with the locus of plasticity. In comparison to the purely postsynaptic case, simulations with presynaptic plasticity presented a smaller variance of the latency shift (Fig. 3B). Potentiation also developed faster with presynaptic expression (Fig. 3F). This can be framed as a consequence of potentiation requiring glutamate release [50], so that in a more reliable synapse, with a high *p* value, there is a greater propensity for potentiation. Conversely, depression was slower with presynaptically expressed plasticity, again because lowered probability of release effectively also led to less plasticity (Fig. 3F).

**Fig 3.**
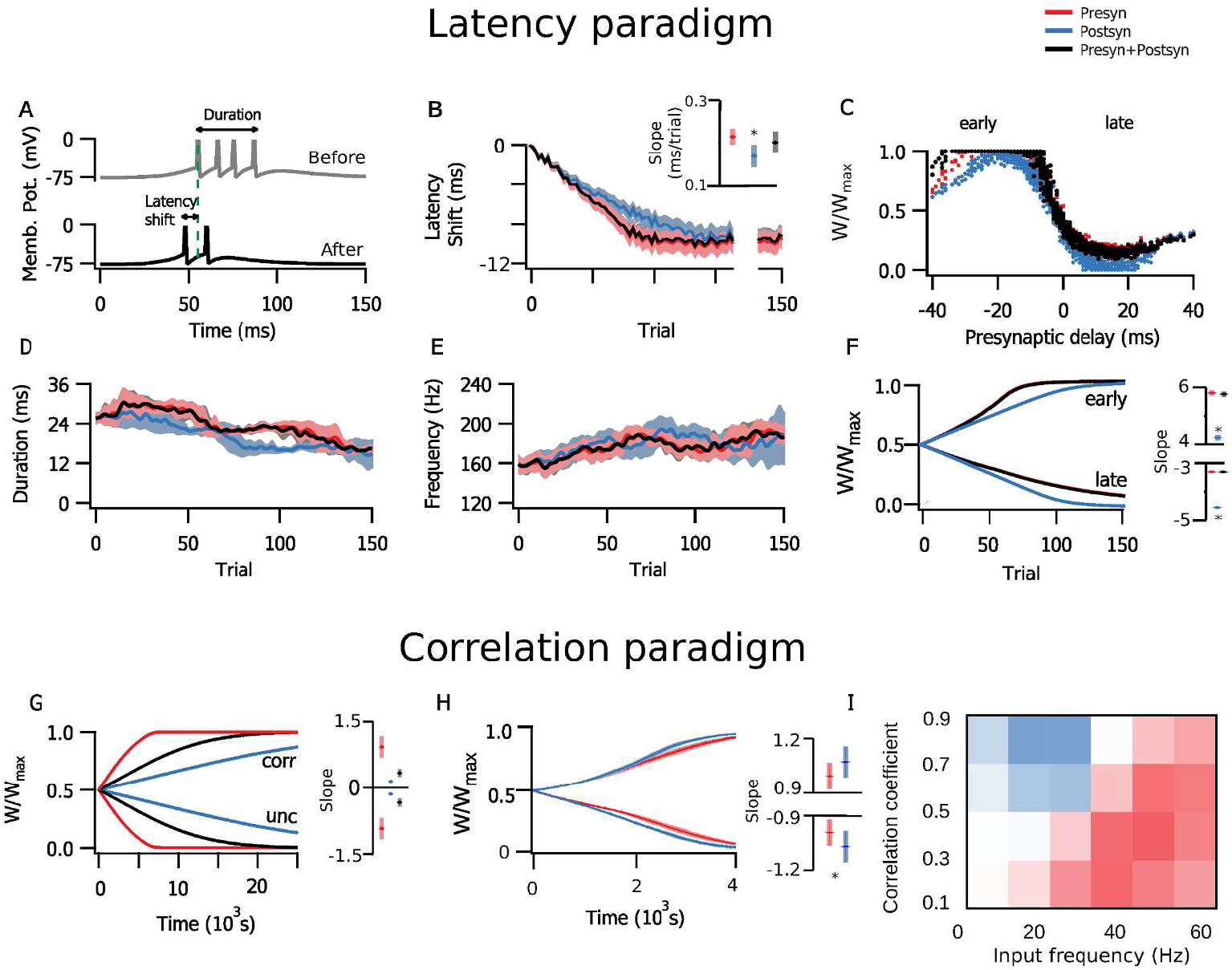
Presynaptically expressed plasticity typically promoted faster learning. (**A-F**) Simulations in the latency paradigm (see Fig. 2A). (**A**) Example traces of the postsynaptic membrane potential before (grey) and after (black) plasticity. Initial latency of response is marked by a green dashed line. (**B**) STDP-mediated learning shortened postsynaptic latency to spike, as previously shown [26]. The graphs here and below are colour-coded: only presynaptic plasticity (red), only postsynaptic plasticity (blue), or both pre- and postsynaptic plasticity (black) are implemented. Note how learning with presynaptic plasticity appears faster. Inset: presynaptic plasticity was faster than postsynaptic plasticity alone (t-test, p-value = 0.008). (**C**) Synaptic weight distribution after 150 trials, normalized and sorted by the fixed presynaptic delay. (**D, E**) Postsynaptic response duration (i.e, the interval between first and last spike in each trial) and the burst frequency did not differ across different expression loci. (**F**) Time evolution of average synaptic weight among early and late presynaptic inputs (i.e., input cells that spiked in the first or the second half of the stimulus) show how post-only expression (blue) is relatively slower. (**G-I**) Simulations in the correlation paradigm (see Fig. 2B). (**G**) Potentiation and depression of the average synaptic weight among correlated inputs was relatively faster in the presynaptic case. (**H**) However, for inputs with a very high correlation (c>0.9), learning was faster with postsynaptic expression. This indicated that which form of plasticity led to faster learning depended on the details of the input firing patterns. (**I**) We explored this finding further in simulations where all inputs were correlated, but half expressed plasticity presynaptically, and the other postsynaptically. The map shows which side potentiated faster, indicating that postsynaptic expression (blue) won out for a relatively small parameter space where input firing frequency was low and and correlations quite high.

Next, we explored the correlation paradigm, in which plasticity selectively potentiates correlated inputs (Fig. 2A,B) ( [45]). Here, all plasticity implementations detected the input correlations. However, presynaptically expressed plasticity generally promoted faster learning, e.g. synaptic weights evolved more rapidly (Fig. 3G), similar to what we found above for the latency paradigm. However, there were exceptions to this general observation — postsynaptic plasticity was faster for strongly correlated input firing at certain input frequencies (Fig. 3H).

We wanted to explore this exception in more detail. One implication of the presynaptic side potentiating faster than the postsynaptic side is that, in a scenario in which there is competition due to e.g. limited resources, inputs with presynaptic plasticity would be expected to overcome inputs with postsynaptic plasticity. To explore this possibility, we ran simulations where all of inputs were correlated, but half of them expressed plasticity only presynaptically, and the other half only postsynaptically. Because of normalization, these two input populations competed, so that one potentiated at the expense of the oher, which depressed. With this approach, we systematically explored the correlation-frequency space. We found that postsynaptic expression won for very highly correlated inputs for sufficiently low input frequencies (Fig. 3I), a scenario that corresponds to fewer inputs spiking synchronously.

### Presynaptic expression modelled as changes in short-term plasticity

We next explored the effects of altering short-term dynamics (see Methods). This adds another aspect of presynaptically expressed plasticity, since short-term plasticity takes into account the history of presynaptic activity. In this scenario, presynaptic changes redistribute synaptic resources used over a limited time period, instead of bringing about an average increase or decrease [16, 41]. Even if the amplitude of an individual EPSP were affected equally by pre- and by postsynaptically expressed plasticity, the total input from a burst would still differ dramatically depending on the site of expression (Fig. 1B).

In the simulations with the timed input configuration, results differed considerably depending on the specific locus of plasticity in the latency configuration. Postsynaptic expression alone provided the largest latency reduction, and also achieved it faster than the other plasticity implementations (Fig. 4A, B). The results of presynaptic expression were also more subtle compared to the mixed setting with both pre- and postsynaptic expression, for which results may vary between extremes according to the ratio of pre- and postsynaptic expression. Effects of postsynaptic plasticity over response duration and intraburst frequency (Figs. 4C and 4D) were also more marked, as expected from a higher integrated input (Fig. 1B). The simulations with both sides changing appeared closer to either the presynaptic case (duration, Fig. 4C) or the postsynaptic case (frequency, Fig. 4D). Here, changes in *p* had a relatively greater influence on response duration, while changes in *q* had greater impact on the response frequency. Nevertheless, synaptic efficacy was still potentiated faster and depressed slower in the presynaptic case (Fig. 4E). This was similar to the above stochastic release implementation of presynaptically expressed plasticity, although it was less pronounced. This means that even if the rate of learning was effectively faster, presynaptic expression affected latency less rapidly than postsynaptic expression did (Fig. 4F).

**Fig 4.**
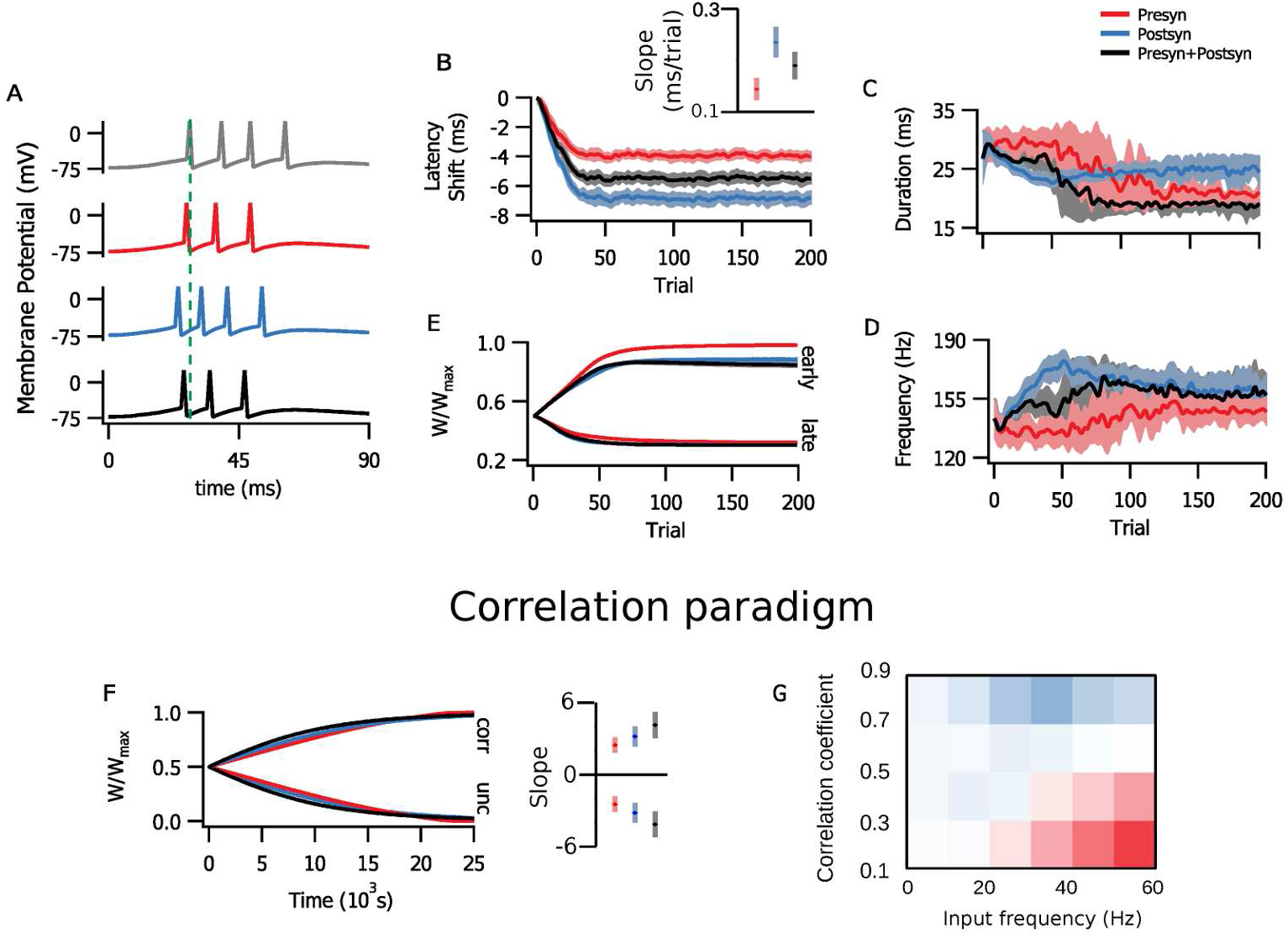
Altering short-term plasticity was less efficient at reducing postsynaptic latency. (**A-E**) Simulations in the latency paradigm (see Fig. 2A) are colour-coded: red denotes presynaptic plasticity alone, blue postsynaptic plasticity alone, and black combined pre- and postsynaptic plasticity. (**A**) Example traces of postsynaptic activity before (grey) and after plasticity (coloured). Initial response latency is illustrated by the vertical dashed line. (**B**) Latency reduction was both faster and more marked for postsynaptic (blue) than for presynaptic (red) or combined (black) plasticity. Inset: The slope of latency reduction was steeper when postsynaptic expression was involved (t-tests: between p and q, p-value < 10^−6^; between p and both, p-value = 0.0008, between q and both p-value = 0.003) (**C**) Combined and presynaptic plasticity reduced response duration more than with postsynaptic expression alone. (**D**) Burst frequency was similarly increased with all three forms of plasticity, although rate change was faster with postsynaptic plasticity. (**E**) Time course of average synaptic weights for early (left) and late (right) inputs. (**F, G**) Simulations in the correlation paradigm (see Fig. 2B) (**F**) Time course of average synaptic weights for correlated (left, “corr”) and uncorrelated (right, “unc”) inputs were largely indistinguishable. (**G**) As with Fig. 3I, the map of competition between input populations with pre- or postsynaptically expressed plasticity indicated a less marked differentiation.

On the other hand, under STP modulation, plasticity rates in the correlated inputs paradigm evolved differently compared to the above stochastic release implementation (Fig. 4G), even though the overall effect on the covariance between pre- and postsynaptic activity was similar (not shown). I the simulations where long-term plasticity affected short-term plasticity, the rate of change was slightly faster with postsynaptic than with presynaptic plasticity. It thus appears that computational advantages could be tailored to a functional task at hand by recruiting pre- or postsynaptic plasticity differentially.

### Comparisons with a biologically tuned model

The above minimalist toy models had the advantage that they provided full control of several key parameters. However, the relevance of the findings for the intact brain were unclear. To address this shortcoming, we explored the biological plausibility in a model [19] (see Methods) that was fitted to long-term synaptic plasticity data obtained from connections between rodent visual cortex layer-5 pyramidal neurons [41, 51, 52]. We could thus to some extent verify whether the results obtained with the minimal models hold in a more complex, data-driven context. We want to clarify upfront that in this model, LTP is expressed both pre- and postsynaptically, whereas LTD is solely postsynaptically expressed. This asymmetry may seem odd, but it is derived from experimental data, and we have previously found that this arrangement provides certain computational advantages [19].

We first explored the latency paradigm (Fig. 2A). To avoid disrupting the parameter tuning, instead of normalising the total synaptic change on each side, we kept the data-derived ratios and blocked either pre- or postsynaptic changes. Even so, we found that both pre- and postsynaptic plasticity components independently led to the shortening of postsynaptic latency (Fig. 5A-C). As with the above simplistic modelling scenarios, postsynaptic changes appeared to affect spike timing more. Thus, when both pre- and postsynaptic plasticity were active, the presence of postsynaptic potentiation further reduced the latency compared to presynaptic plasticity alone (Fig. 5B).

**Fig 5.**
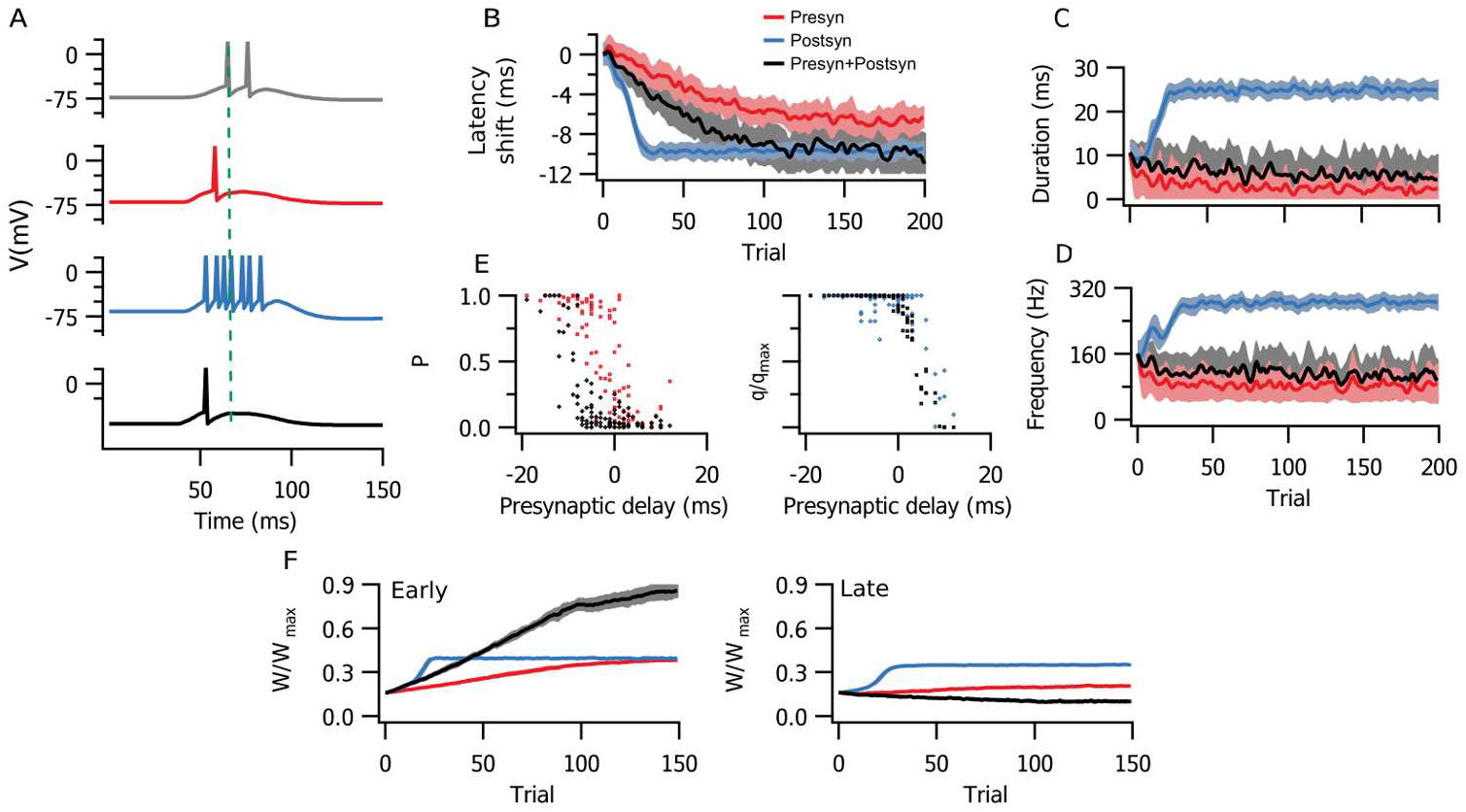
A biologically tuned model verified key findings obtained with minimalist models. (**A**) Sample traces of postynaptic activity before (grey) and after only presynaptic (red), only postsynaptic (blue), or both pre- and postsynaptic learning (black). The initial response latency is indicated by the green dashed line. (**B**) The postsynaptic response latency was shortened by learning, although both faster and more efficiently with postsynaptic learning. (**C**) Distribution of pre- (*p*) and postsynaptic efficacies (*q*) after 200 learning trials. (**D, E**) Changes in duration and burst frequency of postsynaptic activity mirrored those obtained with the stochastic minimalist models (Fig. 4C, D). (**F**) Average synaptic weight of early (left) and late (right) presynaptic inputs evolved in distinct manners, however (compare e.g. Fig. 4).

Following the experimental results [41, 52], postsynaptic plasticity in the tuned model lacked the capacity to depress. As a consequence, postsynaptic plasticity led to inflated postsynaptic frequency and duration when implemented alone (Fig. 5D, E). However, the inclusion of presynaptic LTD was enough to to produce a temporally sharpened response of shorter duration. With postsynaptic plasticity, the dynamics developed faster (Fig. 5F), as result of a positive-feedback loop due to increased postsynaptic firing frequency (compare Fig. 5A).

In the correlation paradigm (Fig. 2A), groups of correlated and uncorrelated inputs clustered (Fig. 6A) without the need for added competition through weight normalization [46, 53]. This only occurred when both pre- and postsynaptic plasticity components were implemented, and was not achieved through other models with physiologically compatible parameters [47].

**Fig 6.**
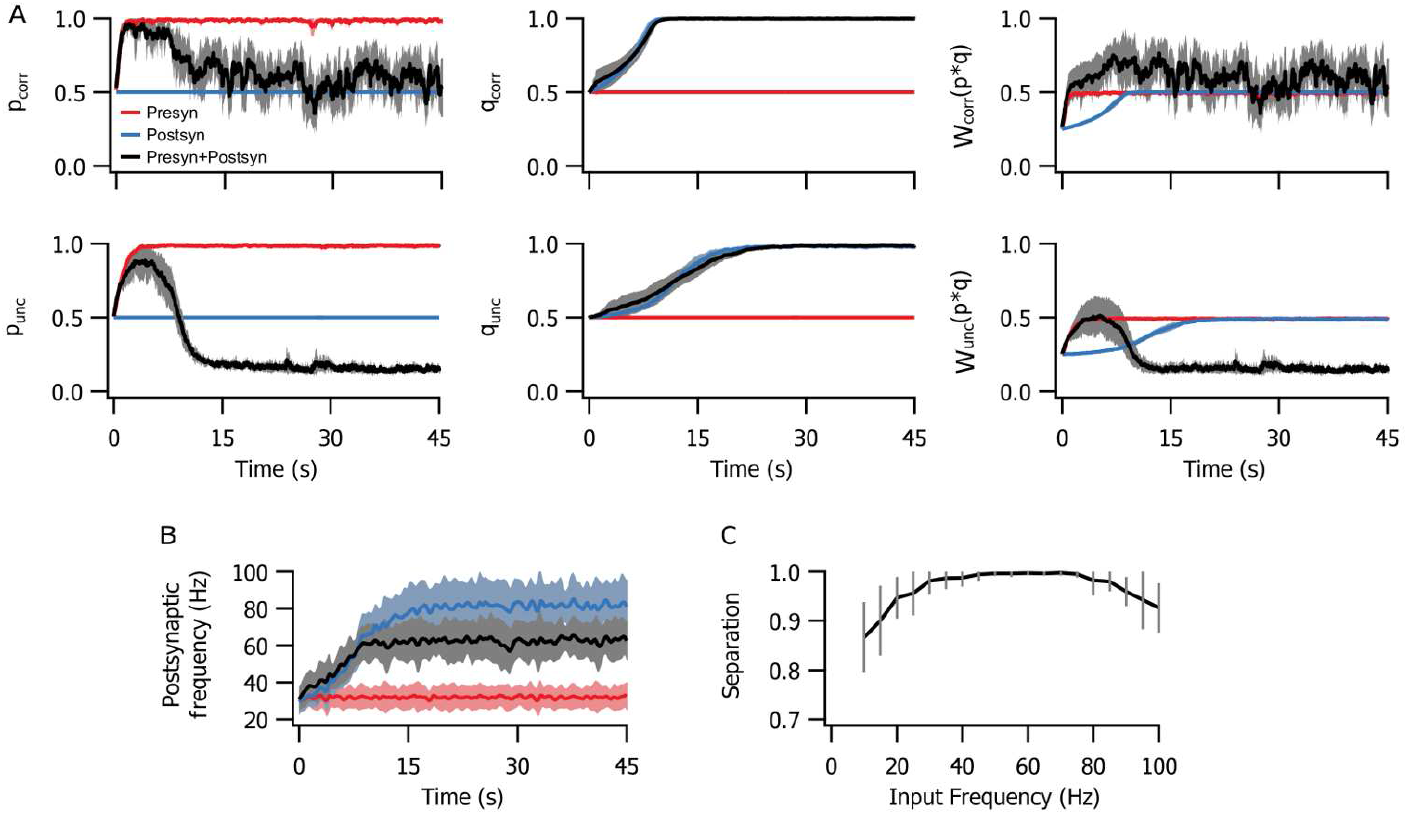
The biologically tuned model clustered inputs with correlated and uncorrelated activity. (**A**) Normalized averages for presynaptic (*p*), postsynaptic and combined pre- and postsynaptic (*W*) plasticity of correlated (corr) and uncorrelated (unc) inputs show that meaningful learning and segregation of inputs occurred when both pre- and postsynaptic learning mechanisms were engaged. (**B**) The postsynaptic spiking frequency increased when postsynaptic plasticity was engaged (blue and black), but not with presynaptic-only learning (red). (**C**) The separation between correlated and uncorrelated inputs was optimal for presynaptic frequencies in the range 50 and 80 Hz.

To better understand the robustness of this property, we quantified the capacity of separation between correlated and uncorrelated populations with a linear separator. It was trained to classify inputs as correlated or uncorrelated according to the average and variance of *p* values (Fig. 6C). The presynaptic frequency range for optimal separation was between 50 and 80 Hz (Fig. 6C). At the other end of the range, it was bounded by the STDP correlation time scale of *τ* = 20 ms (see Methods), meaning interspike intervals longer than 20 ms could not represent the minimal interval of correlation. At the upper end of the range, the high presynaptic frequency yielded overall potentiation that included uncorrelated inputs, limiting the separation from the more potentiated correlated population (see appendix).

In the same way as in the latency paradigm (Fig. 5D), postsynaptic potentiation increased postsynaptic firing rate (Fig. 6B). However, presynaptic plasticity alone produced no such effect. In combination with postsynaptic plasticity, presynaptic plasticity provided a degree of output control, as its introduction helped to maintain a lower postsynaptic firing frequency even as *q* saturated (Fig. 6B).

## Discussion

In recent years, it has become clear that diversity in LTP expression is both ubiquitous and considerable, depending on factors such as animal age, induction protocol, and precise brain region [9–11, 15]. In this work, we explored possible functional properties of either pre- or postsynaptic locus of plasticity expression, and found that even in a single neuron scenario overall dynamics may be affected by it. This is an important feature to be considered, as many theoretical studies have focused on induction but not many in the expression of plasticity. Plasticity has in the typical phenomenological model been implemented by default as a straightforward change in synaptic weight [26, 54, 55], although there are a few notable exceptions [16–18, 56, 57]. In other words, in the absence of better information, a standard assumption has been that that locus of expression does not matter appreciably for the modelling scenario at hand. Our findings challenge this standard assumption, highlighting how it may introduce a bias. For example, over-representation of postsynaptic expression may exaggerate the capacity to learn spike timing (e.g., Figs. 4B, 5B).

We investigated two different learning paradigms, one with differently timed inputs, in which postsynaptic latency to spike was used as a measure of learning (Fig. 2A), and another under constant stimulation, where a subset of inputs were correlated and potentiated together (Fig. 2B). We first worked with simplified conceptual STDP models and later with a more realistic, biologically tuned model in which pre- and postsynaptic components were tuned to connections between neocortical layer 5 pyramidal cells [19].

### Pre- and postsynaptic expression favour different coding schemes

Our study showed that the locus of expression of plasticity determined affinity for different coding schemes. Presynaptic plasticity expressed as the regulation of release probability alone did not result in any differences over average postsynaptic activity measurements compared to postsynaptic expression. However, in the presence of short-term plasticity, presynaptic expression of long-term plasticity had a smaller impact on the spike latency in comparison to postsynaptic expression (e.g., Fig. 4B). This was because, as synaptic response amplitude grew, fewer inputs were needed to evoke a postsynaptic spike. With presynaptic expression, however, the spike still depended on the sum of a larger number of inputs. However, weight changes developed faster with presynaptic plasticity, thereby increasing the speed of learning. This effect, however, was not present in the correlation paradigm, where both pre- and postsynaptically expressed cases performed similarly.

Presynaptically expressed plasticity alone was not ideally suited for rate coding, because it did not impact the average summed input effectively. As a consequence, postsynaptic firing frequency remained relatively unchanged after presynaptically expressed plasticity (e.g., Figs. 4D, 5D, 6B). Presynaptic plasticity thus appeared to act as a limiter or a form of homeostasis for postsynaptic activity, in agreement with previously published interpretations [40]. The flip-side of this stabilizing feature of changes in short-term plasticity [58] is in other words the loss of ability to rate code well. An important cautionary take-home message from this observation is that the default implementation of plasticity as purely postsynaptic may thus lead to an erroneous overestimation of the impact on postsynaptic firing rates.

It is also interesting to think about the role of presynaptic plasticity if it is not very useful in the context of usual ‘coding’ frameworks. Frequently the effect of unreliability of single synapses is considered to be simply of noise or energy economy [59]. However, one can in fact consider this unreliability as a representation of uncertainty over a synaptic weight compared to its optimal value [60, 61]. It would then be plausible to consider pre-synaptic plasticity as an uncertainty tuning over the posterior distribution in a probabilistic inference framework [62].

### A biologically tuned model corroborated the toy model predictions

The same basic properties were observed in the biologically tuned model with simultaneous pre- and postsynaptic plasticity. Learning was dramatically affected by postsynaptic plasticity, while the presynaptic side appeared to act more on the rate of learning and on weight dynamics. It is possible that these results could be modified according to the ratio of pre- versus postsynaptic forms of plasticity, to optimize for the computational task at hand. It is noteworthy that the biologically tuned model was also capable of separating groups of correlated and uncorrelated inputs without the need for a hard competitive mechanism.

### Experimental tests of model predictions

Since it is possible to specifically block pre- or postsynaptic STDP pharmacologically [41, 52], several of our findings related to the locus of expression of plasticity are possible to directly test experimentally. For example, at connections between neocortical layer-5 pyramidal cells, it is possible to block nitric oxide signalling to abolish pre- but not postsynaptic expression of LTP [52]. It is also possible to use GluN2B-specific blockers such as ifenprodil or Ro25-6581 to block presynaptic NMDA receptors necessary for presynaptically expressed LTD without affecting postsynaptic NMDA receptors that are needed for LTP [41, 63]. As a proxy for learning rate, one could explore *in vitro* how blockade of different forms of plasticity expression impacts the number of pairings required for plasticity, or alternatively how the magnitude of plasticity is affected for a given number of pairings [52, 55]. *In vivo*, the impact on cortical receptive fields could similarly be explored. For example, we predict that receptive field discriminability is poorer when presynaptic LTP is abolished by nitric oxide signalling blockade [19].

### Conclusions

Here, we have challenged the standard assumption that modelling synaptic plasticity as a weight change is neutral and unbiased. We found that even in a simple feed-forward scenario, the locus of expression may have considerable impact on learning outcome. We expect that these effect will only be greater in recurrent networks, where presynaptic plasticity at loops and re-entrant pathways will exacerbate the effects of changes in synaptic dynamics due to alterations of the accumulated difference. This additional level of complexity may in particular complicate very large recurrent network models [64, 65].

As our collective understanding of the expression of long-term plasticity has improved, it has become clear that the long-held notion that plasticity is expressed predominantly postsynaptically is erroneous [9–11]. Since presynaptic expression is still relatively poorly studied, our understanding of long-term presynaptic plasticity in health and disease needs to be generally improved [66]. Specifically, our study highlights the need for more detailed modelling of the role of the site of expression.It is clear that it has implications on the relevant form of information coding, be it spike-, rate-based, or probabilistic. In modelling long-term plasticity, correctly implementing changes in weight is thus a matter of gravity.

## Methods

### Neuron model

All of the simulations consisted of one postsynaptic neuron receiving a number of presynaptic Poisson inputs. In the first section, we used a simple leaky integrate-and-fire model defined by

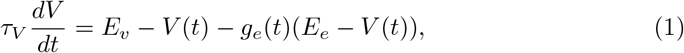

in which the membrane potential *V* decayed exponentially with a time constant of *τ*_*V*_ = 20ms to the resting value of *E*_*v*_ = −74 mV, and the threshold for an action potential was *V*_*th*_ = −54 mV. After each spike it was reset at *V*_0_ = −60 mV with a refractory period of 1 ms.

Inputs were accounted as conductance-based excitatory contributions with reversal potential *E*_*e*_ = 0 mV, amplitude *q*_*j*_, summed after the *l*^*th*^ spike of presynaptic neuron *j*, that decayed exponentially with a time constant of *τ*_*g*_ = 5ms:

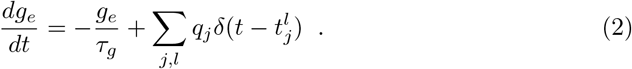

In the the last section, we used the adaptive exponential integrate-and-fire model [67] to reduce unrealistic bursting and to comply with the biological tuning [19]:

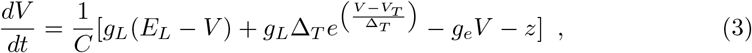

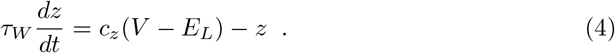

The corresponding parameters for a pyramidal neuron were *C* = 281 pF, *g*_*L*_ = 30 nS, *E*_*L*_ = −70.6 mV, Δ_*T*_ = 2mV, *c* = 4nS, *τ*_*W*_ = 144ms. Spiking threshold was *V*_*T*_ = −50.4 mV, and after each spike *V* was reset to the resting potential *E*_*L*_ while *z* increased by the quantity *b* = 0.0805 nA (as in [67]).

### Stimulation paradigms

The postsynaptic neuron received either one of two stimulus configurations. The first one was based on [26] and is referred to as the Latency Peduction (Fig. 2A). In every 375-ms-long trial, the postsynaptic cell received a volley of Poisson inputs that arrived with a specific delay, normally distributed around a time reference, for each specific presynaptic neuron. Each input lasted for 25 ms with a spiking frequency of 100 Hz. We measured the time to spike of the first postsynaptic spike in response to a bout of stimuli using the mean of the presynaptic delay distribution as a reference point. For clarity, in the Results, curves that represent latency shift, intraburst frequency or burst duration were smoothed using a moving average filter with a window of three points.

The second type of stimulation paradigm was based on [45] and is referred to as the Correlation Paradigm (Fig. 2B). This configuration consisted of continuous Poisson inputs with fixed frequency. However, half of the inputs had correlated fluctuations of activity, with a time window of *τ*_*corr*_ = 20 ms, while the other half was uncorrelated. Correlations were implemente as in [68].

### Additive STDP model

For the majority of the simulations we opted to implement STDP with the simple additive model proposed by Song and Abbott [26]:

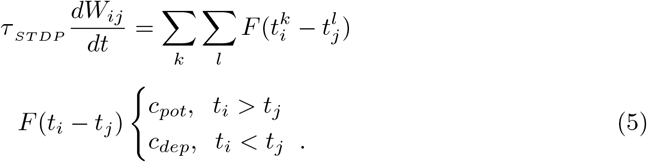

Each increment to the synaptic weights *W*_*ij*_ (since there was only one postsynaptic cell, we consider *W*_*j*_ = *W*_*ij*_ throughout this paper) was computed after a pair of pre- and postsynaptic spikes, and the parameters were set to *τ*_*ST DP*_ = 20ms, *c*_*pot*_ = 0.005, and *c*_*dep*_ = −0.00525. We separated the synaptic weight *W*_*j*_ as a product between pre- and postsynaptic counterparts, probability of release *P*_*j*_ = (0, 1] and quantal amplitude *q*_*j*_ = (0, *q*_*max*_] respectively, so that *W*_*j*_ = *q*_*j*_*P*_*j*_. The probability of release was simulated in two different ways, one equivalent to regulating the probability of stochastic interactions and the other via short-term plasticity.

When the weight convergence rates were compared, we had to ensure that Δ*W* = *W*^*f*^ − *W*^*i*^ per time step was the same for all simulations. Therefore, we normalised the changes so that if only *q* was changed:

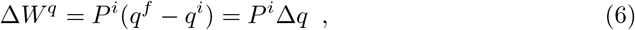

and if only *P* was changed,

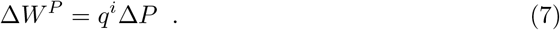

The initial value of all simulations was the same for *P* and *q*, so in these cases Δ*P* = Δ*q* ≡ Δ. This amount was equally divided between *P* and *q* when both were changed simultaneously:

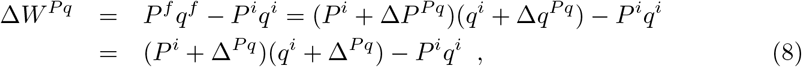

so that

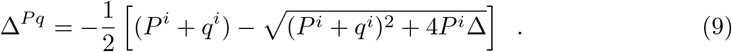

The largest possible change for *P* or *q* separately was Δ_*tot*_ = 1 − *q*^*i*^. To keep the same range of *W* for changing *P* and *q* simultaneously, we limited the maximal values *P* and *q* in this case at 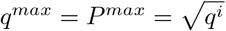.

### Biologically tuned STDP model

We compared the results of the straightforward additive model to a slightly more complex STDP model that acts separately over pre- and postsynaptic factors [19]. Parameters were fitted to experimental data from connections between pyramidal cells from layer 5 of V1 [41, 51, 52]. The equations for pre- and postsynaptic changes followed:

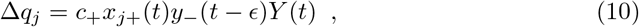

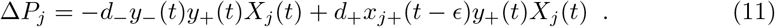

where 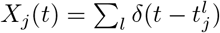 is increased at each spike from the presynaptic neuron *j* and 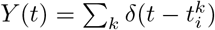 at each spike from the postsynaptic neuron *i*. *ϵ* is to emphasise that Δ*W* was calculated before *x*_*j*+_ and *y*_−_ were updated, upon the arrival of a new spike. *y*_+_ and *y*_−_ are postsynaptic traces,

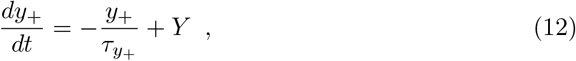

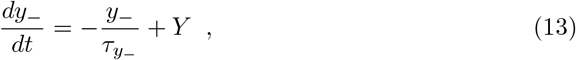

with decay times 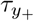 and 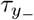 respectively, and *x*_*j*+_ was a presynaptic trace with decay time 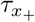:

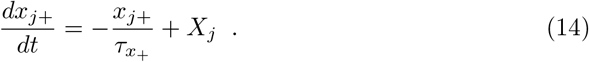

The parameter values were taken from [19]: *d*_−_ = 0.1771, 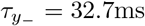, *d*_+_ = 0.15480, *c*_+_ = 0.0618, 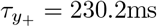 and 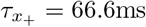. To avoid manipulation of the fitting, weight changes were not normalised in this case.

In the last section, we used a linear least squares separator to classify presynaptic inputs according to synaptic weight average and variance.

### Presynaptic factor

Presynaptic control of the probability of release per stimulus was implemented either as a Markovian process or as short-term plasticity. In the former case, probability (*P*_*j*_) of stochastic neurotransmitter vesicle release followed a binomial distribution. Based on the findings reported by [69], each presynaptic neuron had *N* = 5 release sites that functioned independently. In the second we considered a dynamic modulation of the EPSPs through STP. The probability *P*_*j*_ was decomposed into the product of instantaneous probability of release *p*_*j*_ (*t*) and availability of local resources *r*_*j*_ (*t*), resulting in the following synaptic efficacy:

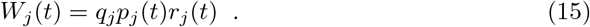

In the latter case, the dynamics of *p*_*j*_(*t*) and *r*_*j*_(*t*) followed the model proposed by Tsodyks and Markram [70]:

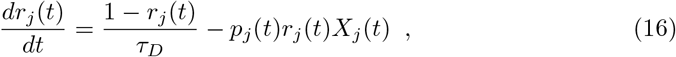

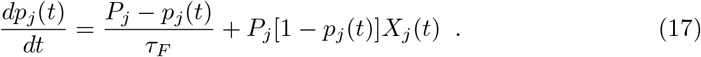

Depression and facilitation time constants, *τ*_*D*_ = 200 ms and *τ*_*F*_ = 50 ms respectively, were chosen as representative values for connections between pyramidal neurons [71]. The resulting short-term plasticity could be either depressing, if *P*_*j*_ > *P*_*C*_, or facilitating, if *P*_*j*_ < *P*_*C*_. For the values of *τ*_*D*_ and *τ*_*F*_ used, *P*_*C*_ ≈ 0.3.

## Supporting information

### S1 Appendix — Rate Model

Using a simple firing rate model with linear response, we were able to illustrate how synaptic plasticity could separate correlated and uncorrelated inputs without competition between the two populations. Considering a neuron receiving independent Poisson inputs with fixed firing rate (pooled into a single average input *I*(*t*)), we found that the system tends to a specific non-zero average value for *P*, denoted *P*∗ below. We converted the biophysically tuned model (eqs. 10 and 11) to a firing rate representation with time-averaged values:

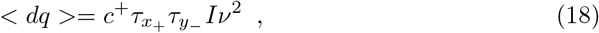

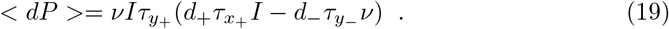

Postsynaptic output *ν* was then considered as a simple firing rate model with linear relation to average input *I*, weighted by average synaptic efficacy (eq. 15):

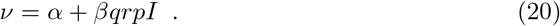

To determine *α* and *β* values that corresponded to the simulated neurons (for fixed values of *q* and *P*), we fitted to data from simulations without plasticity. Since *I* was fixed, we could also consider stationary values for *r*(*t*) and *p*(*t*), 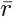 and 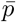, from eqs. 16 and 17:

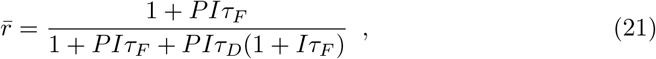

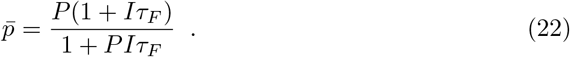

We thus have < *dq* > (*P, q, I*) and < *dP* > (*P, q, I*) in the LTP equations 18 and 19:

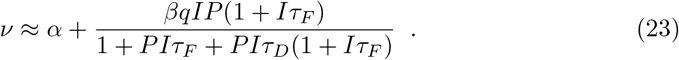

We plotted *dP* × *dq* as a vector field (Fig. 7A), which shows how *P* tended to the specific value *P*∗, which corresponds to the average value of *P* for uncorrelated inputs. Integrate-and-fire simulation averages also converged to this point (black line, Fig. 7A). This is in contrast to correlated inputs, which potentiated more (Fig. 6). The value *P*∗ was relatively stable with frequency, but saturated at a limited frequency value(Fig. 7B), effectivley limiting the range of possible separation between the correlated and the uncorrelated populations.

**Fig 7.**
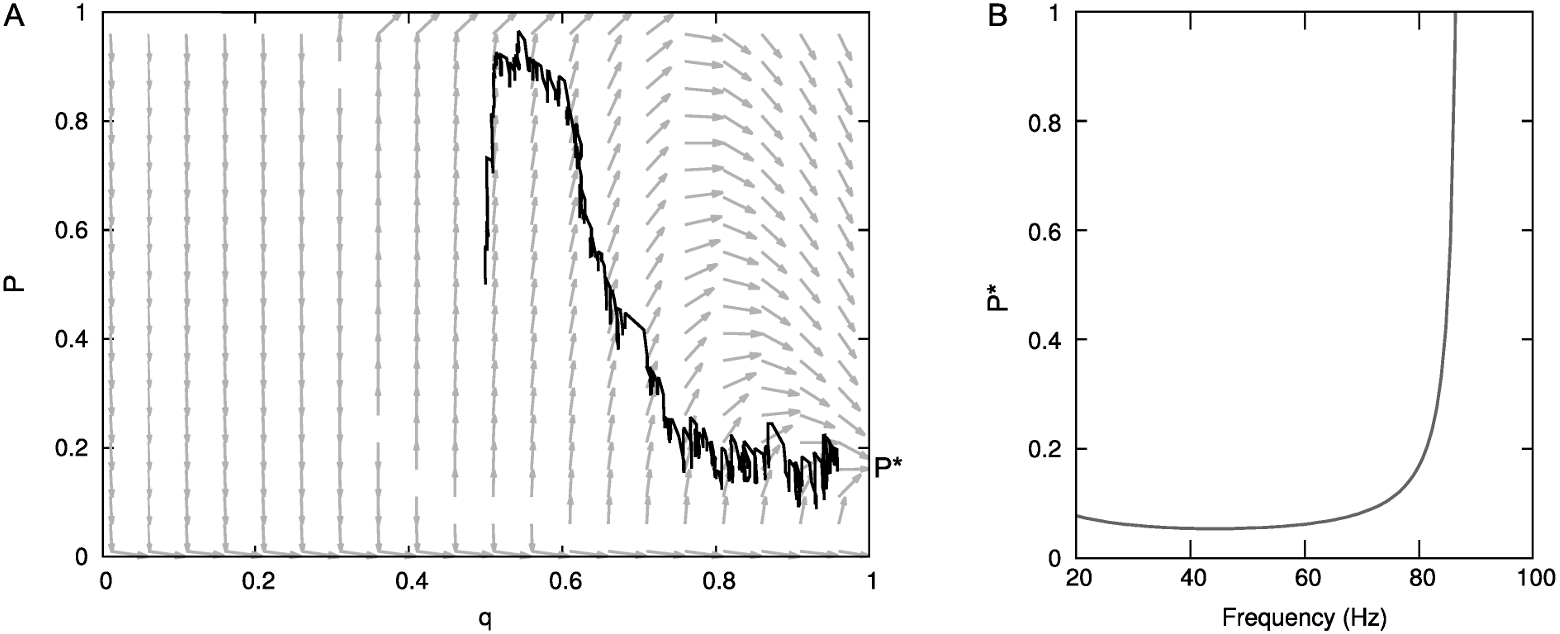
Plasticity separated correlated and uncorrelated inputs up to a limiting frequency. (**A**) Vector field (*p* x *q*) representing the rate model for uncorrelated synaptic inputs only. The black line shows corresponding integrate-and-fire simulation averages for uncorrelated inputs (compare Fig. 6A). Note how both models converge to the same fixed point, indicated with the label *P*∗. (**B**) The point of convergence *P*∗ was relatively stable with respect to presynaptic frequency up to a limiting frequency of around 85 Hz, where it saturated. Since correlated inputs tended to saturate, this shows an effective upper frequency limit to the clustering of correlated and uncorrelated inputs.

## Data availability statement

All files will be available on GitHub.

## Acknowledgements

We thank Alanna Watt and Mark van Rossum for suggestions, help, and useful discussions.

## Author contributions

**Conceptualization:** Beatriz E.P. Mizusaki, Rui P. Costa, P. Jesper Sjöström.

**Formal analysis:** Beatriz E.P. Mizusaki, Sally S.Y. Li.

**Funding acquisition:** Beatriz E.P. Mizusaki, Sally S.Y. Li, Rui P. Costa, P. Jesper Sjöström.

**Investigation:** Beatriz E.P. Mizusaki, Sally S.Y. Li.

**Methodology:** Beatriz E.P. Mizusaki, Rui P. Costa.

**Project administration:** P. Jesper Sjöström.

**Resources:** Rui P. Costa, P. Jesper Sjöström.

**Software:** Beatriz E.P. Mizusaki, Sally S.Y. Li, Rui P. Costa.

**Supervision:** P. Jesper Sjöström.

**Validation:** Beatriz E.P. Mizusaki.

**Visualization:** Beatriz E.P. Mizusaki.

**Writing – original draft:** Beatriz E.P. Mizusaki, Rui P. Costa, P. Jesper Sjöström.

**Writing – review & editing:** Beatriz E.P. Mizusaki, Rui P. Costa, P. Jesper Sjöström.

